# Asymmetric life-history trade-offs shape sex-biased longevity patterns

**DOI:** 10.1101/2025.07.23.665274

**Authors:** Ella Rees-Baylis, Daiping Wang, Xiang-Yi Li Richter, Charlotte de Vries

## Abstract

Sex differences in ageing and lifespan are widespread across taxa, yet their evolutionary causes remain debated. A leading hypothesis suggests these differences are adaptive and driven by sex-specific life-history trade-offs, but formal theoretical support is lacking. To address this, we developed a mathematical model to investigate how such trade-offs shape lifespan evolution in a monogamous mating system. In the model, individuals evolve to optimise a trade-off between reproduction and survival – mediated by mating opportunities in males and offspring production in females. By systematically varying trade-off strengths, we show that either sex can evolve greater longevity, but male-biased longevity evolves under a broader set of conditions – consistent with patterns in monogamous species. This asymmetry arises because female longevity is more constrained: the trade-off between offspring production and survival directly affects the fertility of both sexes. In contrast, the male trade-off for mating opportunities has a weaker indirect effect on female fertility, allowing selection to more readily favour longer male lifespans. We also show that extrinsic density-dependent mortality can disproportionately affect the intrinsically longer-living sex, and obscure the magnitude of this evolved difference. Together, our results provide new theoretical insights into the adaptive bases of sex-biased longevity and highlight the importance of life-history trade-offs in shaping lifespan evolution.

## Introduction

Patterns of sex differences in ageing and lifespan vary widely across and within taxa. For instance, in most mammals, males tend to experience higher mortality rates than females, whereas in birds, sex differences in mortality tend to be smaller but are often female-biased [1–7]. Several non-adaptive, mechanistic explanations have been proposed, including the influence of sex hormones, matrilineal mitochondrial inheritance, and sex chromosome asymmetries [1, 8]. For example, the “unguarded sex chromosome” hypothesis suggests that the heterogametic sex (e.g., male mammals, female birds) may suffer higher mortality due to increased expression of recessive deleterious alleles given only a single X or Z chromosome copy [9, 10]. Furthermore, there is evidence in some species of a “toxic Y (or W)” effect, whereby the higher concentration of repetitive elements on these chromosomes can contribute to reduced longevity as they become misregulated with age [8, 11]. Although the sex determination system correlates with adult sex ratio bias – used as a rough proxy for sex-biased mortality – across multiple taxa [12], empirical support for theories of maladaptive asymmetric inheritance remains mixed [10, 13, 14]. Accordingly, adaptive evolutionary hypotheses have also been proposed to play a major role in shaping sex differences in ageing and longevity.

A predominant idea is that sex differences in lifespan are adaptive, and are shaped by sex-specific selection for different optima in life-history traits [6, 15]. Central to life-history theory is that investment in current reproduction may come at the cost of survival and future reproductive opportunities [16]. Since the sexes often differ in reproductive roles, this trade-off can lead natural selection to favour different optimal strategies in males and females, resulting in sex-specific mortality and longevity patterns [15, 17–19]. Various aspects of reproduction can incur survival costs, including mate acquisition, gamete production, and parental care [20], and as such the nature of this trade-off can differ between the sexes. Females directly produce offspring and often provide more care, both of which can incur survival costs, leading to an evolutionary trade-off between reproduction and survival. For example, avian species with higher annual egg productivity exhibit stronger female-biased mortalities [21] and incubation has been shown to incur physiological and long-term survival costs [22, 23]. Gestation and lactation are particularly energetically costly for mammalian females [24], and more generally across taxa, sex biases in parental care are positively associated with sex biases in adult mortality [5, 25]. In contrast, the primary survival costs of reproduction in males are often linked to competition for mates and the traits arising from this, leading to a trade-off between survival and mating opportunities. Polygynous mating systems – common among mammals – intensify sexual selection in males and are associated with male-biased mortality [2, 26, 27]. Accordingly, the social monogamy typical of most bird species is thought to reduce male-male competition and contribute to higher male survival. This idea is further supported by exceptions across taxa: polygynous bird species tend to show male-biased mortality patterns [5, 21], while the few monogamous mammals often have female-biased mortality [26].

Still, sex-biased mortality patterns in the wild are not only driven by mating systems but emerge from complex interactions between environmental conditions and the costs of sexual selection [28]. Moreover, sexual selection and competition for mating opportunities still persist under monogamy – whether to secure a high-quality partner, obtain and defend a territory, or maintain pair bonds [29, 30]. Thus, whilst mating systems help to explain broad patterns – such as male-biased mortality in predominantly polygynous mammals and female-biased mortality in socially monogamous birds – there remains substantial variation in both the magnitude and direction of sex biases in mortality and lifespan within each mating system that is not yet fully understood.

Furthermore, these current insights have been drawn primarily from empirical studies and meta-analyses, and there remains a lack of formal theoretical predictions to explain how sex-specific life-history trade-offs can shape lifespan evolution. To address this, we constructed a mathematical model that shows how sex-specific survival-reproduction trade-offs influence lifespan evolution under monogamy – a mating system common in birds and also presents in mammals (Fig. 1**a**). In the model, individuals face a trade-off between survival and fecundity: they can evolve to live longer at the cost of reduced annual reproduction, or *vice versa*. The sexes differ only in the mechanism underpinning this trade-off: in males, it arises from mating competition; in females, from investment in offspring production (Fig. 1**b-c**). By systematically varying the strength of these sex-specific trade-offs, we derive predictions for sex differences in life expectancy under different life-history scenarios. Our results provide new theoretical insights into the adaptive basis of sex-biased longevity and highlight the central role of life-history trade-offs in shaping lifespan evolution.

**Fig. 1:**
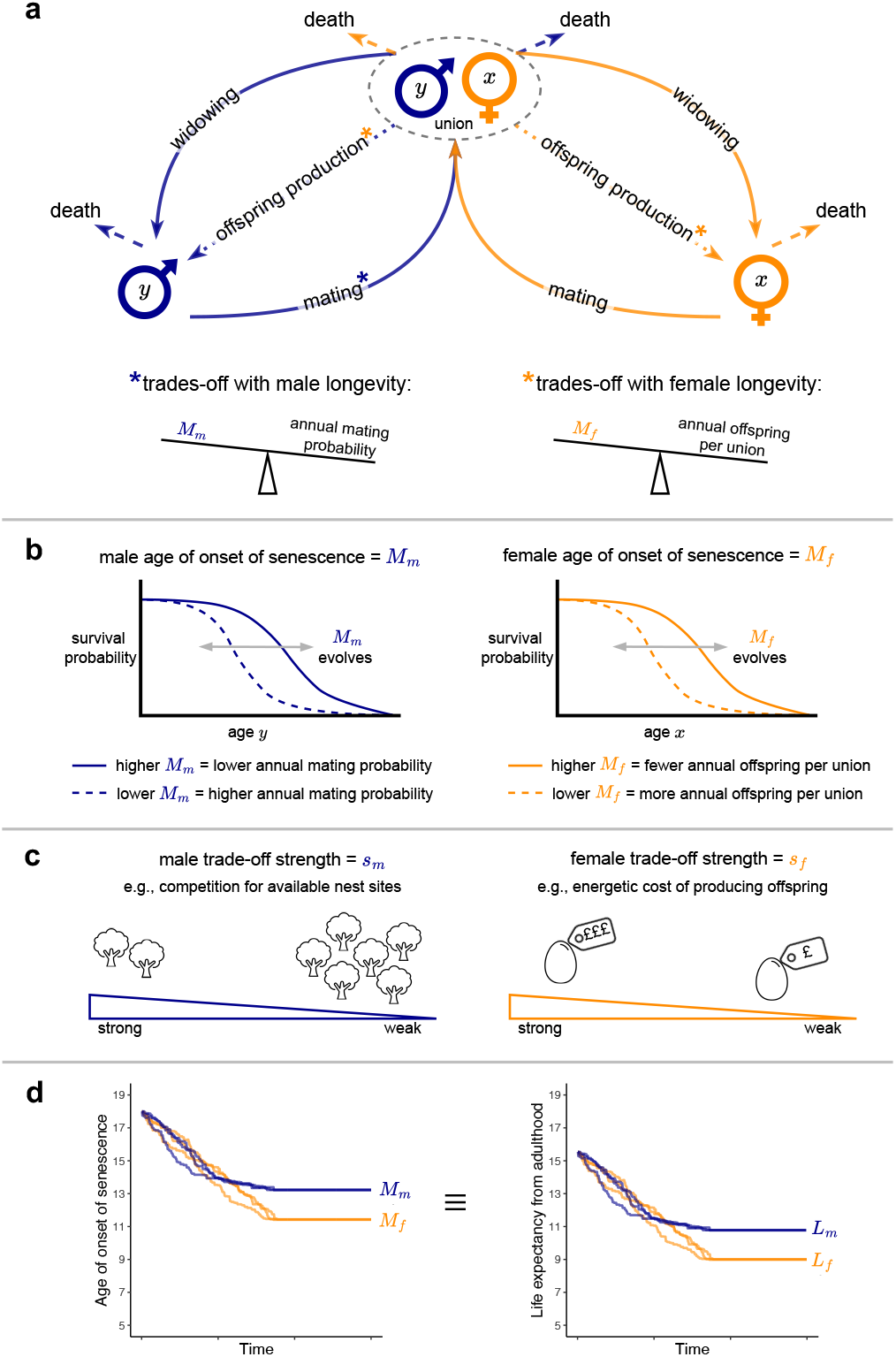
**a** Schematic overview of the model. Blue and orange arrows represent male and female transitions, respectively. The population consists of single males (aged *y*), single females (aged *x*), and monogamous unions of one male and one female. Individuals can age from 1 to 20, but are removed upon death. Singles must mate to form unions, which can then produce offspring (*i*.*e*., new single males and females in age class 1). If one union member dies, the surviving partner returns to the singles pool and may re-mate. A male’s annual mating probability trades off with his longevity, while the number of offspring produced per union per year trades off with female longevity. **b** Sex-specific survival-reproduction trade-offs. *Left* : male age of onset of senescence (*M*_*m*_) evolves to balance survival and mating opportunities – later onset increases life expectancy but reduces annual mating probability. *Right* : female age of onset of senescence (*M*_*f*_) evolves to balance survival and fecundity – later onset increases life expectancy but reduces the annual number of offspring per union. **c** Examples of trade-off strengths. *Left* : the male trade-off may arise from competition for nest sites. When sites are abundant, mating is less costly and *s*_*m*_ is weak; when sites are scarce, intensified competition imposes greater survival costs, strengthening the trade-off. *Right* : the female trade-off may stem from energetic demands of reproduction. When offspring are inexpensive to produce, *s*_*f*_ is weak; when offspring require high energy investment, survival is more compromised, resulting in a stronger trade-off. **d** Example evolutionary trajectories of the male and female ages of onset of senescence. For clarity and comparisons with empirical data, results are plotted as the corresponding equilibrium life expectancies from adulthood (a y-axis transformation; see Methods for details).

## Results

For all model results, we examine the evolved equilibrium values of the sex-specific ages of onset of senescence – that is, the age at which senescence significantly reduces survival. The male age of onset (*M*_*m*_) evolves to optimise the trade-off between survival and annual mating probability, while the female age of onset (*M*_*f*_) evolves to balance survival with the annual number of offspring produced per monogamous union (Fig. 1**b-c**). These traits evolve toward a stable evolutionary equilibrium that is robust to initial trait values (Fig. 1**d**; Fig. S1). We investigated how the strength of the male and female trade-offs (*s*_*m*_ and *s*_*f*_, respectively) influence the evolved equilibrium ages of onset of senescence. For easier interpretation and comparability with empirical data, we present the results in terms of male and female life expectancies at adulthood (*L*_*m*_ and *L*_*f*_, respectively), since this is a standard metric to report (sex-specific) longevity (Fig. 1**d**; see Methods for details).

### Males outlive females under a wider range of trade-off strengths

In general, stronger survival-reproduction trade-offs lead to the evolution of shorter lifespans in both sexes (Fig. 2; males **a1**-**2**, females **b1**-**2**). Consequently, the sex with the weaker trade-off typically outlives the one with the stronger trade-off, meaning either males or females may have the longer lifespan depending on trade-off assumptions (Fig. 2 **c1 & c2**). However, when trade-off strengths are equal – or when the male trade-off is only slightly stronger than the females’ – males tend to evolve to marginally longer lifespans, resulting in male-biased longevity across a broader range of trade-off strengths.

**Fig. 2:**
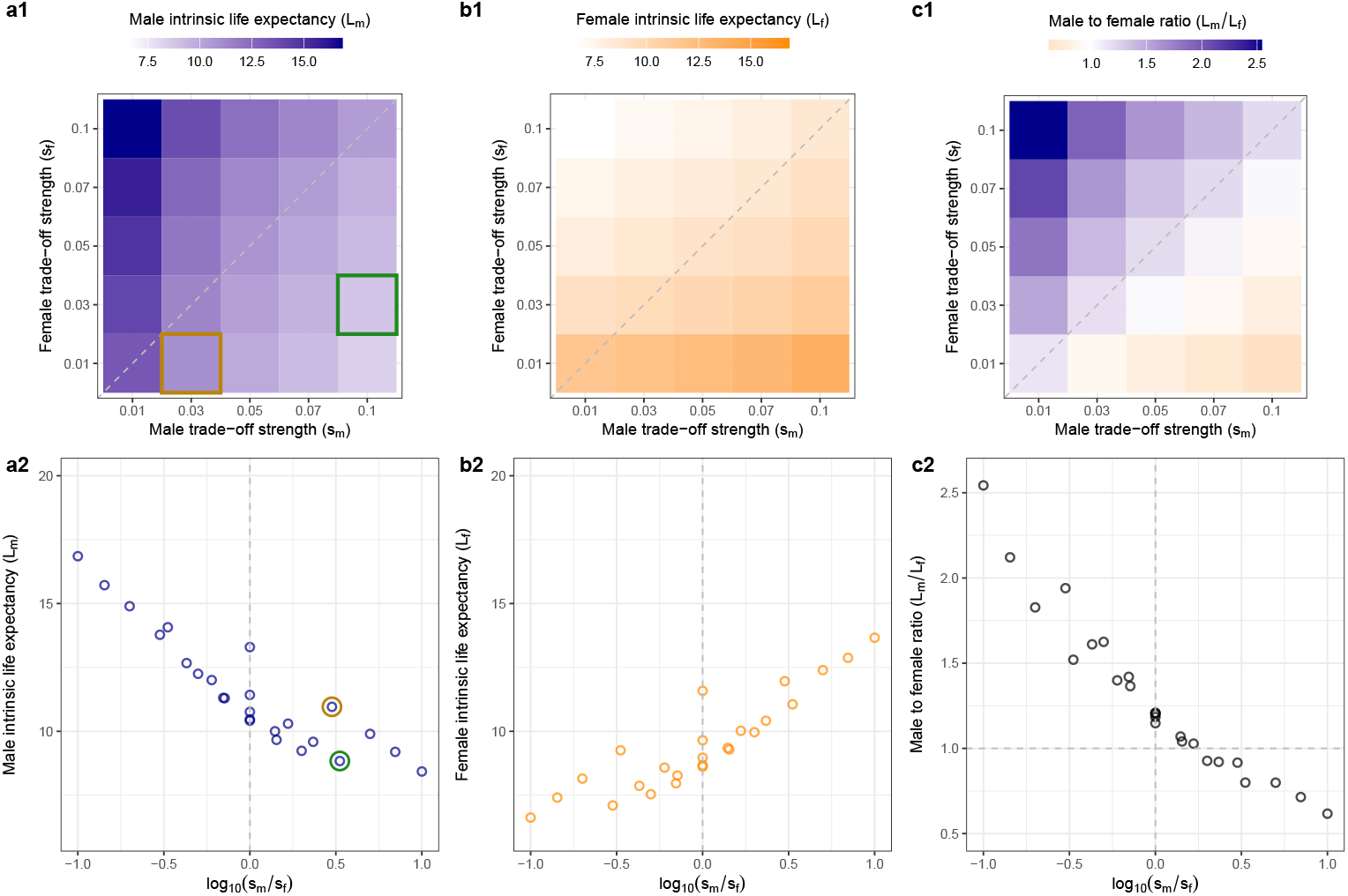
Heatmaps (**a1-c1**) and scatter plots (**a2-c2**) showing the effects of sex-specific survival-reproduction trade-offs on evolved life expectancy. Heatmaps show (**a1**) male intrinsic life expectancy (*L*_*m*_), (**b1**) female intrinsic life expectancy (*L*_*f*_), and (**c1**) the ratio of male to female intrinsic life expectancy (*L*_*m*_/*L*_*f*_) across a range of male (*s*_*m*_) and female (*s*_*f*_) trade-off strengths. Diagonal dashed lines indicate equal trade-off strength between the sexes (*s*_*m*_ = *s*_*f*_). Scatter plots (**a2-c2**) present the same data as a function of the relative trade-off strength between sexes (log_10_(*s*_*m*_/*s*_*f*_)). Vertical dashed lines indicate equal trade-off strengths. In **c2**, the horizontal dashed line denotes parity in life expectancy between the sexes (*L*_*m*_ = *L*_*f*_); points above this line indicate male-biased longevity, while those below indicate female-biased longevity. Apparent “noise” in the scatter plots arises because different absolute trade-off strength combinations yield similar ratios (e.g., log_10_(0.1/0.03) = 0.52 ≈ log_10_(0.03/0.01) = 0.47; highlighted in green and yellow in **a1-2**).

We next explored a broader parameter space to asses how additional reproductive constraints influence the optimisation of survival-reproduction trade-offs and, in turn, the evolution of sex differences in life expectancy. Specifically, we varied two parameters: the maximum annual fecundity (the number of offspring a union can produce per year, which is diminished by the female trade-off), and the mating efficiency (the proportion of the mating pool that will mate to form unions under balanced population sex ratios; [31]).

Figure 3 shows that the main pattern – where either sex may evolve to be longer-lived, but males tend to outlive females across a wider range of trade-off strengths – remains robust across different values of maximum annual fecundity. Consistent with pace-of-life theory, both sexes evolve shorter life expectancies as the potential annual number of offspring produced increases. Additionally, higher maximum annual fecundity reduces both the magnitude of sex biases in life expectancy and the parameter space in which males outlive females.

**Fig. 3:**
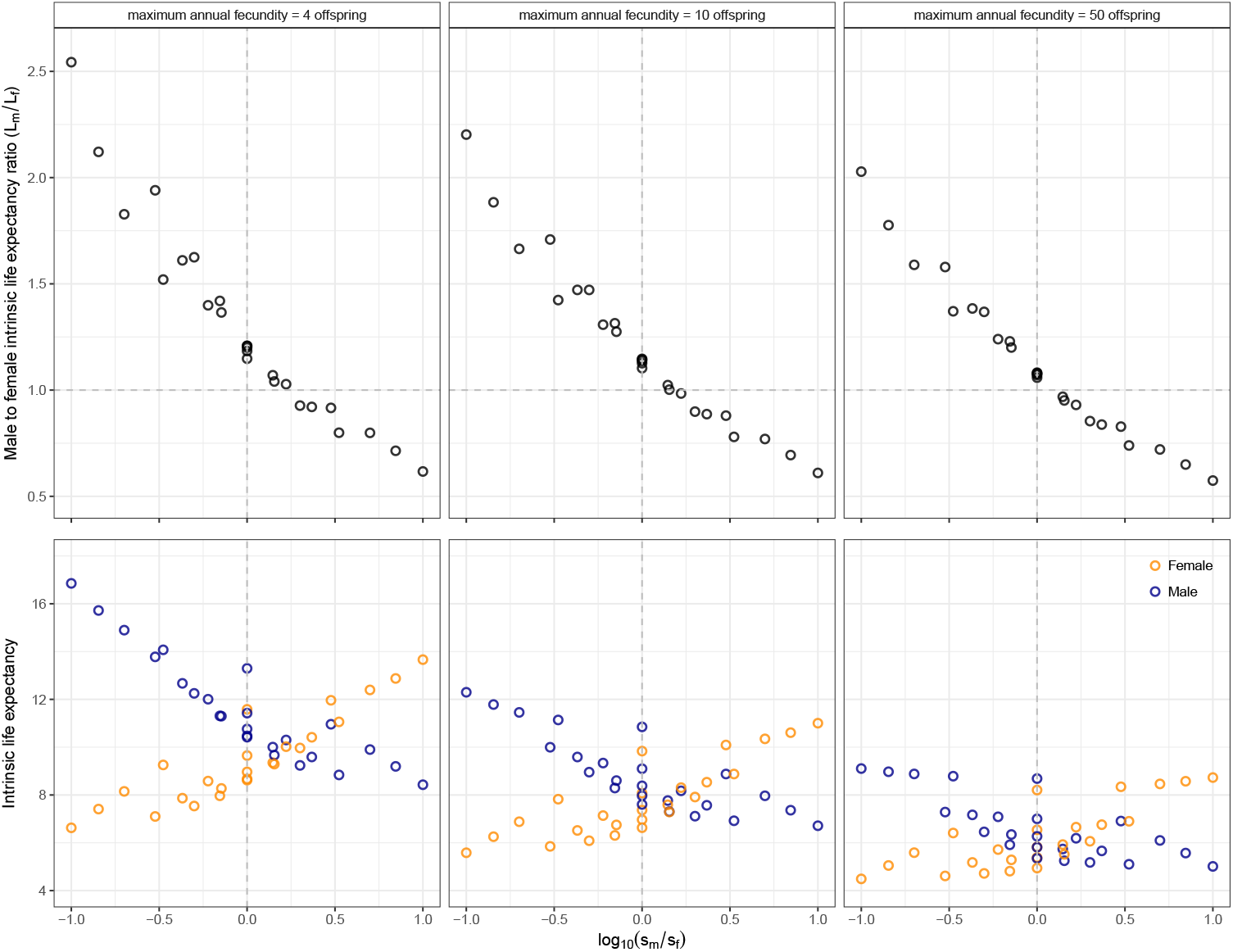
*Top*: male-to-female life expectancy ratios across the relative trade-off strength between sexes (log_10_(*s*_*m*_*/s*_*f*_)) under varying maximum annual fecundity values. *Bottom*: the corresponding evolved life expectancies in each sex. Vertical dashed lines indicate equal trade-off strength between the sexes (*s*_*m*_ = *s*_*f*_). In the top row, horizontal dashed lines represent parity in life expectancy (*L*_*m*_ = *L*_*f*_); points above this line indicate male-biased longevity, while those below indicate female-biased longevity.

Our main results are also robust to changes in mating efficiency (Fig. S2). However, increased mating efficiency slightly reduces life expectancy in both sexes. This suggests that when mating is less efficient, there is a stronger selective advantage to prolonged lifespan – allowing individuals to reproduce over more years – rather than having a higher annual reproductive output over a shorter life.

### Female determination of union reproduction constrains female longevity

Why do males outlive females across a broader range of trade-off strength scenarios in our model? Importantly, a trade-off in one sex can influence not only the fitness of individuals of that sex, but also fitness of the opposite sex. In our model, these cross-sex effects differ between the male and female trade-offs. We assume that female longevity trades off with the number of offspring produced per year per mating union. As a result, female longevity directly and strongly affects both female and male fertility – since males depend on the reproductive output of their monogamous female partners (Fig. 4**a**). In contrast, male longevity primarily influences male fertility, with only a weaker, indirect on female fertility through mating dynamics (Fig. 4**b**). This asymmetry places stronger constraints on the evolution of female longevity, making it more costly for females to live longer. Consequently, males evolve to outlive females under a wider range of trade-off scenarios.

**Fig. 4:**
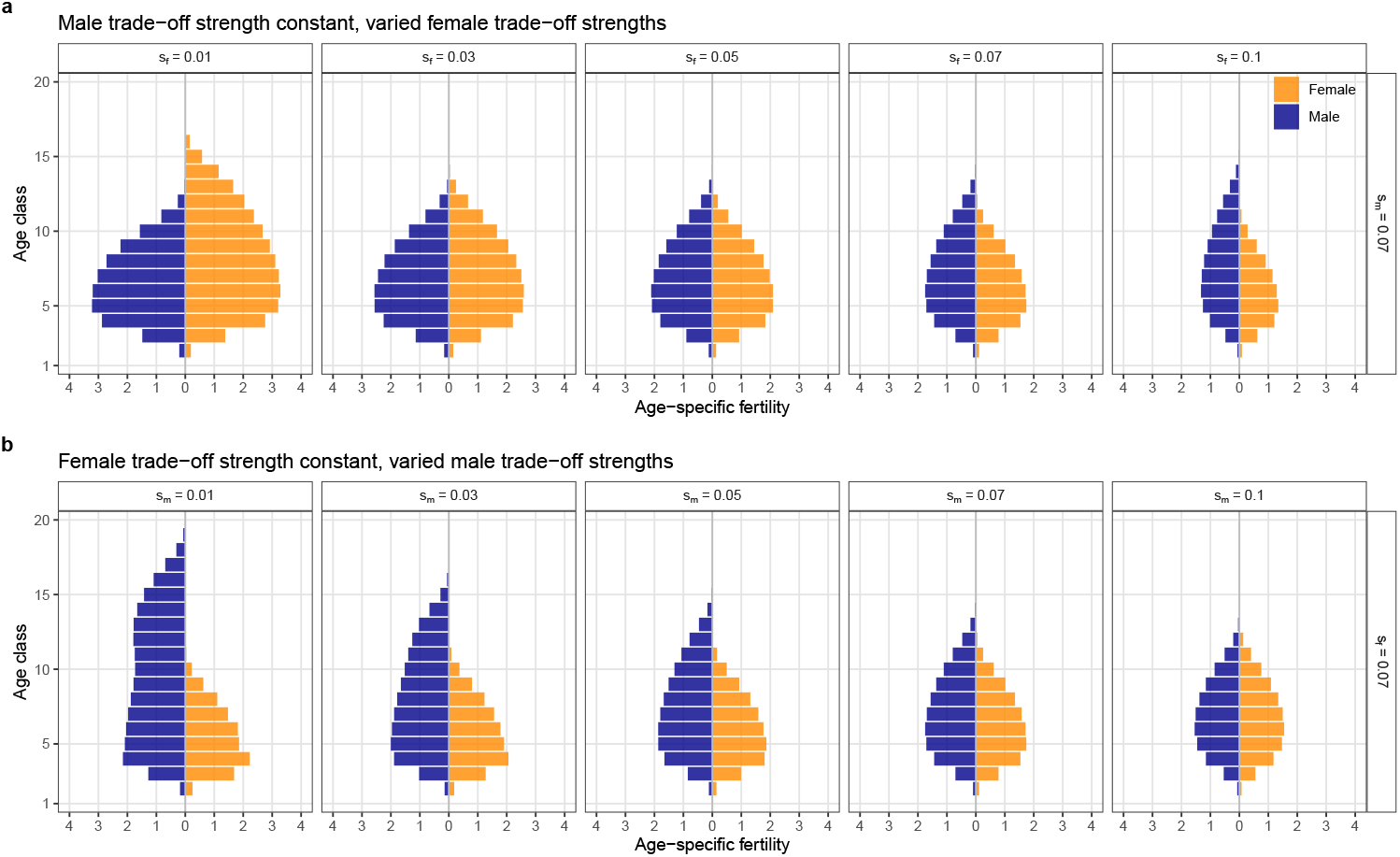
**a** Age-specific fertility (average number of offspring produced at a given age, conditioned on survival to that age) for each sex across a range of female trade-off strengths but a fixed male trade-off strength of *s*_*m*_ = 0.07. **b** Age-specific fertility for each sex across a range of male trade-off strengths but a fixed female trade-off strength of *s*_*f*_ = 0.07. See Fig. S3 for age-specific fertility patterns across all combinations of trade-off strengths.

In addition, the male trade-off also influences the relative timing of first reproduction. When males evolve longer lifespans, their annual mating probability is decreased, delaying their age at first reproduction relative to females. Conversely, shorter male lifespans lead to earlier breeding compared to females. Thus, the male trade-off strength indirectly determines which sex achieves higher early-life reproductive output – typically the sex with the shorter lifespan (Fig. 4; Fig. S3).

### Density regulation can conceal intrinsic sex differences in survival

The results so far have shown sex-biases in longevity as a result of evolved differences in senescence – that is, differences in intrinsic survival with age. However, in natural populations, individuals are also subject to extrinsic sources of mortality. Density-dependent mortality, for example due to competition for limited resources, is particularly common since few populations grow exponentially. To account for this, we compare *intrinsic* life expectancies (based solely on senescence) with *realised* life expectancies, which incorporate both intrinsic and density-dependent survival.

As a simple illustrative case, we assume that density-dependent mortality acts equally across all age classes and both sexes – for instance, under uniform resource limitation where all individuals require the same amount of resources. Under this assumption, our main result still holds: males outlive females under a broader range of trade-off conditions. This is because equal density dependence does not alter the intrinsic ages of onset of senescence that evolve [32–34]. However, the magnitude of sex differences in realised life expectancy is substantially reduced compared to the intrinsic differences alone (Fig. 5**a**). Although both sexes experience the same additional density-dependent mortality per time step, the longer-lived sex – alive for more time steps – experiences a stronger compound effect of this mortality. This compounding effect diminishes the observed sex difference in life expectancy. Importantly, this implies that sex asymmetries in survival-reproduction trade-offs can result in large intrinsic survival differences between the sexes, but that empirically these differences may become only marginally detectable due to the effects of density regulation. Sex differences in life expectancy are often subtle across monogamous species (Fig. 5**b-c**), highlighting the difficulty of disentangling evolved, intrinsic survival differences from those imposed by ecological constraints such as density-dependent mortality.

**Fig. 5:**
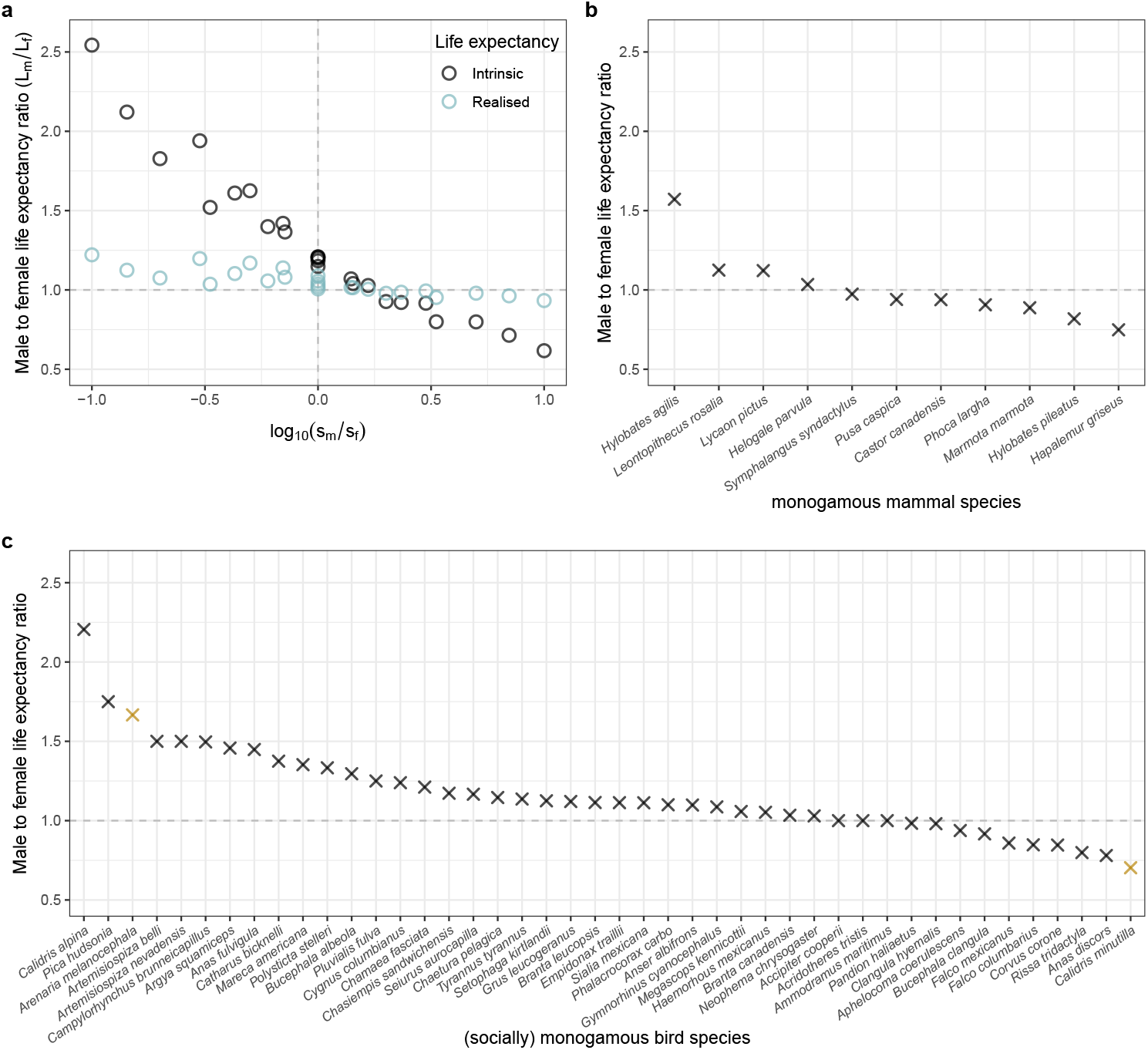
**a** Model-predicted male-to-female life expectancy ratios across the range of relative trade-off strengths between the sexes (log_10_(*s*_*m*_*/s*_*f*_)). Black points indicate intrinsic life expectancy ratios (based solely on senescence); blue points show the realised life expectancy ratios (incorporating both intrinsic and density-dependent survival). **b** Empirically measured male-to-female life expectancy ratios from *n* = 11 monogamous mammal species. **c** Empirically measured male-to-female life expectancy ratios from *n* = 44 socially monogamous bird species. For species in which data was available for more than one population, the point represents the average ratio across populations. Yellow points indicate species mentioned as examples in the Discussion. See Methods for details on empirical data sourcing.

## Discussion

We here presented a mathematical model providing a theoretical framework for understanding how sex-specific trade-offs between survival and reproductive success can drive differences in lifespan between the sexes under monogamous mating systems. By systematically varying the strength of these trade-offs – between survival and annual mating probability in males, and survival and annual union offspring production in females – we demonstrate that either sex may evolve to live longer, generally depending on which faces the weaker trade-off. This result offers a potential explanation for the observed variability in sex-biased longevity among (socially) monogamous species: while slight female-biased mortality is common, male-biased patterns also occur (e.g., [2, 5]). Our model suggests that this variation could stem, at least in part, from sex differences in life-history trade-offs.

For example, in the monogamous least sandpiper (*Calidris minutilla*), females tend to outlive males (Fig. 5**c**). Males in this species provide extensive parental care, including incubation and often sole responsibility for chick rearing after females depart early [35]. In our model, this could reflect a very weak trade-off between survival and offspring production in females – or a strong one in males – parameter regions where female longevity is expected to exceed male longevity. Conversely, in the monogamous black turnstone (*Arenaria melanocephala*), males tend to live longer than females (Fig. 5**c**). These birds exhibit strong mate fidelity and return annually to the same territory to mate with the same partner [36], which in our model may correspond to a weak trade-off between survival and mating opportunities in males, supporting greater male longevity.

These examples are, however, illustrative. While empirical support for early-late trade-offs across a wide range of species exists [37], many other factors beyond intrinsic trade-offs – such as ecological constraints and environmental variability – also affect sex- or species-specific ageing and lifespan patterns 17, 38, 39]. For instance, although strong sexual selection in males (akin to a strong male trade-off strength in our model) is predicted to result in shorter lifespans and/or faster ageing in males, sexual selection proxies such as sexual size dimorphism or testes size often show inconsistent relationships with sex-biased longevity and ageing rates [15, 40, 41]. This challenges the centrality of life-history trade-offs n shaping sex differences in longevity and ageing and highlights the need for targeted empirical studies – such as manipulations of reproductive investment or survival costs – as well as further theoretical work that incorporate different assumptions.

Although our model assumes monogamy, it opens a path towards deeper theoretical understanding of how mating systems can shape sex-specific longevity. Notably, our male trade-off strength parameter (*s*_*m*_) can be viewed as a proxy for the survival costs of male-male competition for reproduction, which are likely more extreme in polygynous or promiscuous mating systems. In our model, stronger male trade-off strengths generally result in males living shorter than females, consistent with patterns observed in polygynous species [2, 15, 27]. However, explicit predictions for different mating systems will require models that incorporate these structures directly, ideally within a comparative framework.

The model also revealed that males tend to evolve longer lifespans than females across a broader range of trade-off scenarios. This arises from an asymmetry in how longevity affects fitness in the two sexes, shaping the model’s demographic structure. Specifically, we assume that only female longevity determines annual offspring production of a mating pair – reflecting the common modelling assumption of female demographic dominance (or partial dominance, since female fecundity in our model is still reduced when males are the limiting sex). Although it is sometimes argued that female demographic dominance should not affect evolutionary outcomes in diploid organisms, since both sexes contribute equally on average to reproduction, theoretical work has shown otherwise [42–44]. Our model highlights another non-trivial consequence: female survival has a direct and stronger influence on population-level reproduction, imposing tighter evolutionary constraints on female lifespan. In contrast, in our model male longevity affects access to mates but not offspring production directly, allowing selection to more readily favour longer male lifespans – especially when the survival-mating trade-off is weak. Crucially, our results suggest female-biased mortality as the default outcome under female determination of pair reproduction, with the direction of bias reversible depending on the strength of male reproductive costs. Other factors that reduce the extent of female demographic dominance may also shift outcomes; for example, species with more extensive male parental care tend to show greater male-biased mortality [5].

In this context, sex-role reversed species provide valuable test cases for understanding sex-specific longevity. In the sequentially polyandrous barred buttonquail (*Turnix suscitator*), males assume primary care-giving and reproductive investment roles, yet females still fundamentally produce each clutch. Here, males live on average 1.7 times longer than females [45] – indicating that the survival costs of producing offspring in females can outweigh those of raising them in males. Nevertheless, while we implemented sex-specific trade-offs based on two common hypotheses – the survival costs of sexual selection in males and of offspring production or care in females – more general models are needed to capture the diversity of sex-specific life histories and to better understand how the extent of female demographic dominance influences outcomes.

Our model also incorporates density-dependent mortality in a simplified, uniform manner affecting all individuals equally. Under this assumption, density regulation can greatly diminish the observed magnitude of evolved intrinsic sex differences in lifespan: the longer-lived sex accrues a greater exposure to density-dependent mortality, offsetting some of its survival advantage from the weaker intrinsic trade-off. Whilst this serves as an important conceptual point, in nature, density dependence is unlikely to act uniformly across individuals. Extrinsic mortality risks – such as predation, disease, or competition – may vary by sex, age, or condition. Furthermore, the sex roles that we assumed here to result in different intrinsic survival-reproduction trade-offs between the sexes will likely also result in differential exposures to extrinsic hazards. For example, competition for mating opportunities in males may not only incur intrinsic survival costs but also increase exposure to predation [46] and increase parasite risk [47]. Our model also assumes that all individuals are of equal quality, but individual heterogeneity can obscure trade-offs [48, 49], with high-quality individuals often exhibiting both higher fecundity and longer lifespans compared to lower-quality ones (e.g., [50, 51]). Exploring how such sex-, condition-, and context-dependent mortality influences the evolution of sexual dimorphism in ageing and lifespan remains an important direction for future theoretical and empirical work.

Another important consideration in the evolution of sex biases in longevity is sexual conflict [6, 20, 52]. Our model assumes no intra-locus sexual conflict – that is, no genetic constraint arising from shared longevity-affecting loci under sexually antagonistic selection – allowing the sexes to genetically evolve independently towards their respective optima and potentially resulting in strong lifespan dimorphism. In reality, shared genetic architecture could limit the extent to which each sex can independently optimise its strategy [6, 53]. For instance, artificial selection experiments in *Callosobruchus maculatus* beetles have shown that males live longer and females live shorter than their sex-specific optima as a result of genetic correlations for lifespan between the sexes [54]. Such correlations would likely reduce the magnitude of sex differences in lifespan as predicted by our model. Nonetheless, our model does highlight the role of inter-locus sexual conflict: fitness interests of one sex can impose costs on the other through reproductive interactions [20]. For instance, in our model, lower male mating effort can reduce fecundity of females if males become limiting as a consequence. In this sense, our model underscores that sexual dimorphism in ageing and longevity does not emerge solely from independent sex-specific optimisation, but also from the interactive consequences of reproductive strategies.

Finally, we modelled the evolution of sex-specific life expectancies assuming fixed, non-dimorphic ageing rates. While much empirical data also focuses on longevity metrics, sex differences in longevity do not necessarily imply sex differences in ageing rates, indicating that much remains unknown about the evolution of sex-specific ageing and senescence of other traits [4, 15, 55]. More longitudinal studies in wild populations are needed to better understand the evolution of sex-specific ageing. Moreover, much of our understanding of early-late trade-offs comes from laboratory studies, often measured only in females - presumably since female reproductive effort is easier to quantify [37]. Not only should future empirical work aim to test for survival-reproduction trade-offs in males, but ideally both sexes in the same wild population. Our model, like work on sexual conflict, underscores the importance of reproductive interactions in shaping sex differences in longevity, highlighting the need for life-history data from both sexes.

In conclusion, our model provides a general first theoretical framework for how sex-specific trade-offs between survival and reproduction can shape sex-biased lifespan evolution under monogamy. It identifies key mechanisms – asymmetric trade-off strengths, demographic influence, and sexual interactions – that may help explain variation in the magnitude and direction of sex-biased longevity across taxa. For instance, male-biased longevity may evolve as a result of female determination of pair reproduction, unless males have strong reproductive costs that can shift the bias towards longer female longevity. Future theoretical work should explore the effects of explicit mating system structures, sexual conflict, individual heterogeneity, and non-uniform extrinsic mortality. At the same time, more longitudinal data from wild populations – capturing longevity metrics, ageing rates, and other life-history traits in both sexes – are essential to deepen our understanding of sex-specific ageing and lifespan evolution.

## Methods

### Model overview

We constructed a discrete-time, two-sex, age-structured matrix population model [56]. The population is divided into single males, single females, and monogamous unions formed by one male and one female (Fig. 1**a**). Senescence is modelled using the standard Gompertz hazard function, where individuals experience an exponentially increasing risk of death with age. We assume that this mortality trades off with reproduction, such that each sex can evolve to live longer at the cost of reduced reproductive output, and *vice versa*. In males, this trade-off operates through their annual probability of joining a union (*i*.*e*., mating opportunities); in females, it acts through the annual number of offspring produced per union (Fig. 1**b-c**). This is the only sex difference in the model, so the labels “male” and “female” are arbitrary: for example, the trade-offs could be reversed in systems with female-female competition or male-biased parental care. We used an adaptive dynamics framework to allow the age of onset of senescence – the age at which senescence substantially decreases survival – to evolve in each sex, with evolution proceeding until they reach a stable evolutionary equilibrium (Fig. 1**d**; Fig. S1). We varied the strength of the sex-specific trade-offs between survival and reproduction to observe the effects this has on the equilibrium longevities that evolve in each sex. All model parameters and their default values are summarised in Table 1.

**Table 1:** Model parameters and their (range of) values used, unless stated otherwise.

| Parameter | Meaning | Value(s) | Equation(s) |
| --- | --- | --- | --- |
| $\omega$ | Number of age classes | 20 | |
| $b$ | Rate of increase of mortality with age | 0.06 | 1 |
| $M_m$ | Male age of onset of senescence | Evolving | 1,7 |
| $M_f$ | Female age of onset of senescence | Evolving | 1,8,9 |
| $g$ | Maturation function shaping parameter | 5.0 | 4 |
| $h$ | Maturation function shaping parameter | 2.5 | 4 |
| $\eta$ | Mating efficiency | 0.8 | 6 |
| $s_m$ | Male survival-reproduction trade-off strength | [0.01, 0.03, 0.05, 0.07, 0.1] | 7 |
| $s_f$ | Female survival-reproduction trade-off strength | [0.01, 0.03, 0.05, 0.07, 0.1] | 8,9 |
| $k$ | Maximum annual union fecundity | 4 | 8,9 |
| $r$ | Primary sex ratio (proportion male) | 0.5 | 8,9 |
| $\sigma$ | Mutational step size | 0.1 | |
| $v$ | Strength of density dependence | 0.0001 | 17,20 |

### Survival

Individuals age from 1 to a maximum age *ω*, advancing to the next age class if they survive the discrete time step. The probability that an individual of sex *i* (*m* for males and *f* for females) survives from age class *x* to *x* + 1 is given by

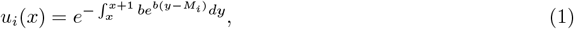

where *b* is the rate of increase of mortality with age, and *M*_*i*_ is the age of onset of senescence for individuals of sex *i. M*_*i*_ represents the age at which senescence (*i*.*e*., intrinsic mortality) begins to strongly limit survival, and corresponds to the modal age at death in the absence of extrinsic mortality. The term 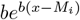 is the Gompertz hazard rate, a widely used model of senescence that assumes an exponentially increasing risk of mortality with age [57]. Taking the exponential of the negative integral of this function over the interval [*x, x* + 1] yields the discrete survival probability for that time step [58]. All individuals die upon reaching the maximum age class, such that *u*_*i*_(*ω*) = 0. The age of onset of senescence for males and females (*M*_*m*_ and *M*_*f*_, respectively) are the evolving traits in the model. Their equilibrium values represent how sex-specific life histories evolve to optimise the trade-off between survival and reproduction.

### Union formation

To reproduce, single males and females must first form monogamous mating unions. Union formation is modelled using a flexible, frequency-dependent *minharmonic mating function* [31].

We first calculate the size of the mating pool. The number of single males available to mate is given by:

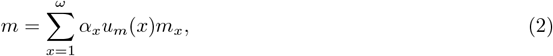

and likewise, the number of single available females is:

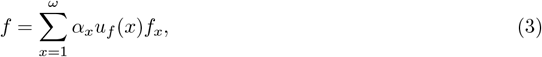

where *m*_*x*_ and *f*_*x*_ are the numbers of single males and females in age class *x*, respectively. The term *α*_*x*_ represents the age-specific reduction in mating probability due to immaturity, modelled as:

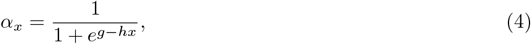

where *g* and *h* determine the shape of this logistic maturation function. As age increases, *α*_*x*_ approaches 1, describing a gradual transition from reproductive immaturity to maturity.

The total size of the mating pool is:

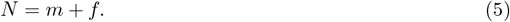

By summing over all age classes, we assume that mating is random with respect to age among individuals that are sufficiently mature enough to mate (as defined by equation (4)).

Following [31], the expected number of unions that form each time step is given by the minharmonic mating function:

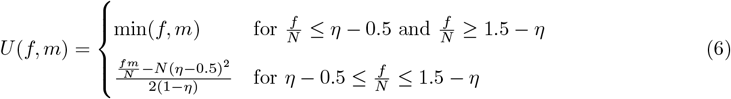

where *η* is the mating efficiency; the probability of mating at balanced operational sex ratios. For example, *η* = 1 means all available individuals form unions at balanced sex ratios, whereas *η* = 0.8 implies that only 80% of the mating pool does so, despite the balanced sex ratio. This mating function ensures that the number of unions formed is limited by the rarer sex under skewed sex ratios, and approximates the harmonic mean between sexes under balanced sex ratios.

The number of unions formed between a male of age *y* and a female of age *x* is then given by:

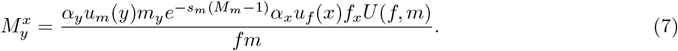

This expression is the product of three terms: the per-pair probability of union formation 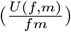 the number of mating-ready females of age 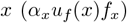 and the number of mating-ready males of age *y*, modified by their evolved mating effort 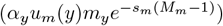. The term 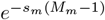 captures the males’ reduction in annual mating probability, determined by the strength of the male survival-reproduction trade-off (*s*_*m*_) and the male’s evolved age of onset of senescence (*M*_*m*_). Thus, males can evolve longer lifespans (higher *M*_*m*_), but at the cost of reduced annual mating success (Fig. 1**b**).

### Reproduction

Each time step, all unions produce offspring, with output independent of the ages of the parents within each union. The number of male offspring produced per union per time step (*i*.*e*., the number of new single males entering age class 1) is given by:

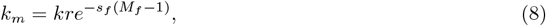

and likewise, the number of female offspring (*i*.*e*., new single females of age class 1) is:

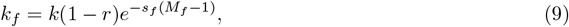

where *k* is maximum annual fecundity of a union, *r* is the primary sex ratio (proportion of male offspring), and 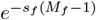 is the reduction in maximum annual fecundity, according to the strength of the female survival-reproduction trade-off (*s*_*f*_) and the current evolved female age of onset of senescence (*M*_*f*_). Thus, females can evolve longer lifespans (higher *M*_*f*_), but at the cost of reduced annual reproductive output per union (Fig. 1**b**).

Although not explicitly implemented, one can interpret individuals in the model as haploid, carrying two unlinked loci with sex-specifically expressed genes: one affecting the male trait (*M*_*m*_) and one affecting the female trait (*M*_*f*_). Offspring inherit one allele at each locus, randomly contributed by the two parents. In the resident population, this inheritance has no phenotypic consequence, as all individuals share the same genotype (*M*_*f*_, *M*_*m*_).

### Union survival & dissolution

A mating union persists to the next time step only if both the male and female members survive. The probability that a union between a male of age *y* and a female of age *x* survives is given by:

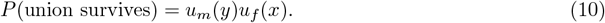

If either partner dies, the surviving individual is returned to the singles pool and becomes eligible to form a new union in the following time step. The probability that a female of age *x* becomes widowed (*i*.*e*., her male partner of age *y* dies) and returns to the singles pool is:

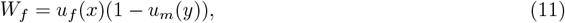

and likewise, the probability that a male of age *y* becomes widowed due to the death of a female partner of age *x* is:

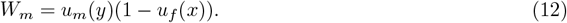

### Matrix construction & population projection

The population at time *t* is represented by the array:

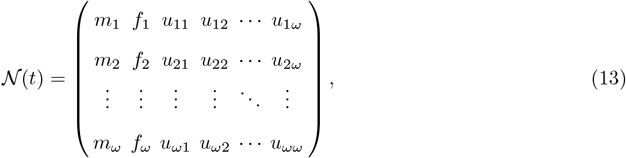

where *m*_*x*_ and *f*_*x*_ denote the number of single males and females of age *x*, respectively, and *u*_*xy*_ represents the number of unions between females of age *x* and males of age *y*. This two-dimensional array is transformed into a one-dimensional population state vector using the vec operator, which stacks the columns of the array:

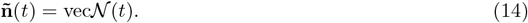

The resulting population state vector consists of three main blocks:

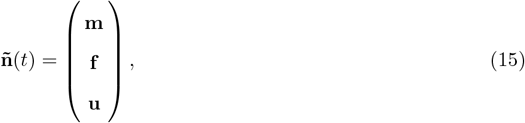

where

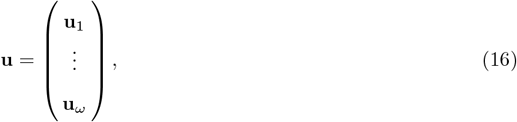

and **u**_*x*_ contains all unions involving a female of age *x* with males of any age.

The population vector is projected forward in time using a frequency-dependent projection matrix:

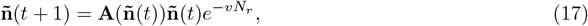

Where 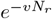 is a density-dependent term that regulates population growth. Here, *v* is the strength of density dependence (*i*.*e*., determining population size at carrying capacity), and *N*_*r*_ is the total population size, calculated as:

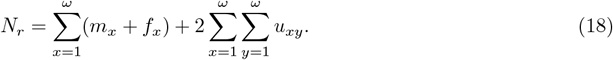

By scaling all elements of the projection matrix with this density-dependent term, we assume a simplified scenario in which density regulation affects both sexes and all age classes equally.

Given the three blocks in the population vector (equation (15)), the projection matrix has a 3 × 3 structure:

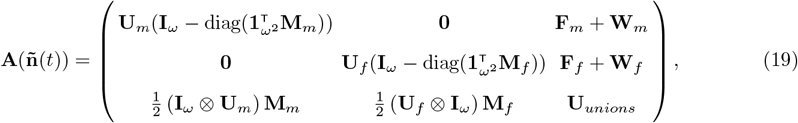

where:

- **U**_*m*_ and **U**_*f*_ contain the age-specific survival rates of males and females, respectively.
- **M**_*m*_ and **M**_*f*_ contain the age-specific mating probabilities (*i*.*e*., transition into unions) for males and females, respectively.
- **F**_*m*_ and **F**_*f*_ contain the reproductive rates of unions to produce sons and daughters, respectively.
- **W**_*m*_ and **W**_*f*_ contain the widowing rates of unions to produce single adult male and females, respectively.
- **U**_*unions*_ contains the age-combination-specific survival rates of unions.

Together, these sub-matrices represent the demographic processes described previously - survival, union formation, reproduction, and union survival and dissolution. See the Supplementary Information for details of their explicit structures and construction of the projection matrix.

### Evolution via adaptive dynamics

We used an adaptive dynamics framework to model the evolution of the male and female ages of onset of senescence (*M*_*m*_ and *M*_*f*_, respectively), simulating these dynamics to obtain numerical solutions. At each time step, we assessed whether a rare mutant – with a slightly altered age on onset of senescence (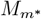 or 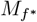) – could invade a resident population at equilibrium. This process was repeated until the trait values converged to a numerical equilibrium, representing uninvadable singular strategies (Fig. S1). The resident population was projected through time using the frequency-dependent resident population projection matrix (equation (17)). Over time, the population converges to a stable equilibrium distribution 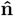, with a long-term growth rate given by the dominant right eigenvalue of 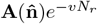, denoted as *λ* [59]. Since we assume density regulation occurs (implemented using the term 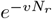), *λ* converges to 1.0.

To evaluate mutant invasion, we constructed a mutant population projection matrix **A**^∗^, which retains the structure of the resident matrix (equation (19)) with the exception that there are two types of unions: resident male-mutant female and mutant male-resident female ([60]; see the Supplementary Information for details). Because mutants are rare, we assume mutant-mutant unions do not occur. The mutant projection matrix **A**^∗^ was calculated using a mutated age of onset of senescence value, 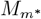 or 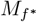, generated by changing the resident value (*M*_*m*_ or *M*_*f*_) with a small random value drawn from a Gaussian distribution with mean *µ* = 0 and standard deviation *σ* (the mutational step size).

The implicit model assumption that individuals are haploid, carrying both *M*_*m*_ and *M*_*f*_ genes but expressing only the one corresponding to their sex, is relevant for mutant dynamics. We assume that mutations arise sequentially, alternating between loci. When a mutation arises at the *M*_*m*_ locus, mutant females are phenotypically identical to resident females because they express the wild-type *M*_*f*_, but they carry the mutant 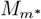 allele and can pass it to their offspring with a probability of 0.5. Similarly, when a mutation arises at the *M*_*f*_ locus, mutant males are phenotypically identical to resident males because they express the wild-type *M*_*m*_, but carry the mutant 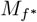 allele and can pass it to their offspring.

The growth rate of the rare mutant was calculated in the environment defined by the resident equilibrium distribution, 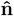. Accordingly, the mutant growth rate, *λ*^∗^, is the dominant right eigenvalue of 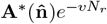, where *N*_*r*_ still represents the population size of residents (equation (18)) and not mutants, since we assume rare mutants do not influence population dynamics [60]. Invasion fitness was evaluated by comparing *λ*^∗^ to the resident *λ*: a mutant can successfully invade and replace the resident if *λ*^∗^ *> λ* (*i*.*e*., if *λ*^∗^ *>* 1, given the implementation of density regulation).

In summary, evolutionary dynamics are simulated by iteratively introducing small mutations to either *M*_*m*_ or *M*_*f*_, calculating the respective growth rates *λ* and *λ*^∗^, and updating the trait value if the mutant invades successfully (*i*.*e*., if *λ*^∗^ *> λ*). This process is repeated until the trait values converge to a numerical equilibrium, corresponding to a singular strategy. To account for minor numerical variation in calculations, we ran three replicate simulations per scenario and report the mean evolved values of *M*_*m*_ and *M*_*f*_ across replicates.

### Calculation of life expectancies

In the model, the evolving traits of interest are the male and female ages of onset of senescence (*M*_*m*_ and *M*_*f*_, respectively). However, to aid interpretation of model results and to allow for comparison with empirical data, we instead report the male and female life expectancies at adulthood (*L*_*m*_ and *L*_*f*_, respectively). To achieve this, we use the age-specific survival matrices for males and females (**U**_*m*_ and **U**_*f*_, respectively), which contain age-specific survival probabilities along the sub-diagonal (Eq. 1; see Supplementary Information for details). These survival matrices are used to calculate the fundamental matrix **N**_*i*_ for each sex *i*, where each element *n*_*xy*_ represents the average time an individual of age *y* can expect to spend in age class *x* [59]:

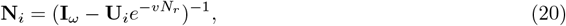

where **I**_*ω*_ is an *ω* × *ω* identity matrix with 1’s along the diagonal and 0’s elsewhere. Summing each column of **N**_*i*_ yields an *ω* × 1 vector of life expectancies: the average time until death for a sex *i* individual of each age class. We define life expectancy from adulthood as the second element of this vector – corresponding to age class 2 – the first age class with non-zero reproduction in our model.

We distinguish between two types of life expectancy: *intrinsic* and *realised*. Intrinsic life expectancy reflects how long an individual is expected to live based on senescence only, excluding any extrinsic mortality. For this, we set *v* = 0 in equation (20) to remove the added density-dependent mortality from the life expectancy calculation. Realised life expectancy (shown only in Fig. 5) reflects how long individuals can be expected to live on average given their intrinsic survival and the additional mortality as a result of density regulation. For this, we set *v* = 0.0001 in equation (20), the default parameter value used in simulations (Table 1).

### Empirical data

The empirical data presented in Figure 4**b-c** was sourced from [10] (bird and mammal data), the “Birds of the World” database [61] (bird data), and [28] (mammal data). For each source, we used all species that are classified as strictly (socially) monogamous and for which average life expectancies from adulthood of both sexes was available. Data in [10] and [61] contain sex-specific life expectancies, but not a mating system classification. For species from these datasets, we classified bird mating systems using the definitions in [62] and for mammals using the data from the “Animal Diversity Web” database [63] if available for the species, and appended this to the life expectancy data. These criteria resulted in sex-specific life expectancies for *n* = 44 (socially) monogamous bird species and *n* = 11 monogamous mammal species.

## Data availability

The model was constructed and run using MATLAB v23.2.0 [64]. Data output from the model was subsequently plotted in R v4.4.0 [65] using the packages *tidyverse* [66], *patchwork* [67], and *ggpubr* [68]. All model code, plotting scripts, and files with extracted empirical data are available on request.

## Acknowledgements

We would like to thank Colin Olito and Stefano Giaimo for discussions about the model, and Hanna Kokko for input on modelling assumptions. We are also grateful to the IT Service Office at the University of Bern for providing access to the high performance computing cluster UBELIX, where simulations were run. We also acknowledge funding from the Swiss National Science Foundation (211549) to X-Y.L.R., and from the CAS Pioneer Hundred Talents program and the Institute of Zoology, Chinese Academy of Sciences (2023 and 2024) to D.W.

## Supplementary Information

### Evolutionary equilibria

We model the evolution of the male and female ages of onset of senescence (or for presentation of results, the life expectancies at adulthood) via an adaptive dynamics framework. The male and female life expectancies therefore evolve to equilibrium values that are evolutionary stable strategies (ESSs): a rare mutant with a different male or female life expectancy value cannot invade the resident population at equilibrium. Therefore, these equilibrium values are reached regardless of the initial trait values (Fig. S1).

**Fig. S1:**
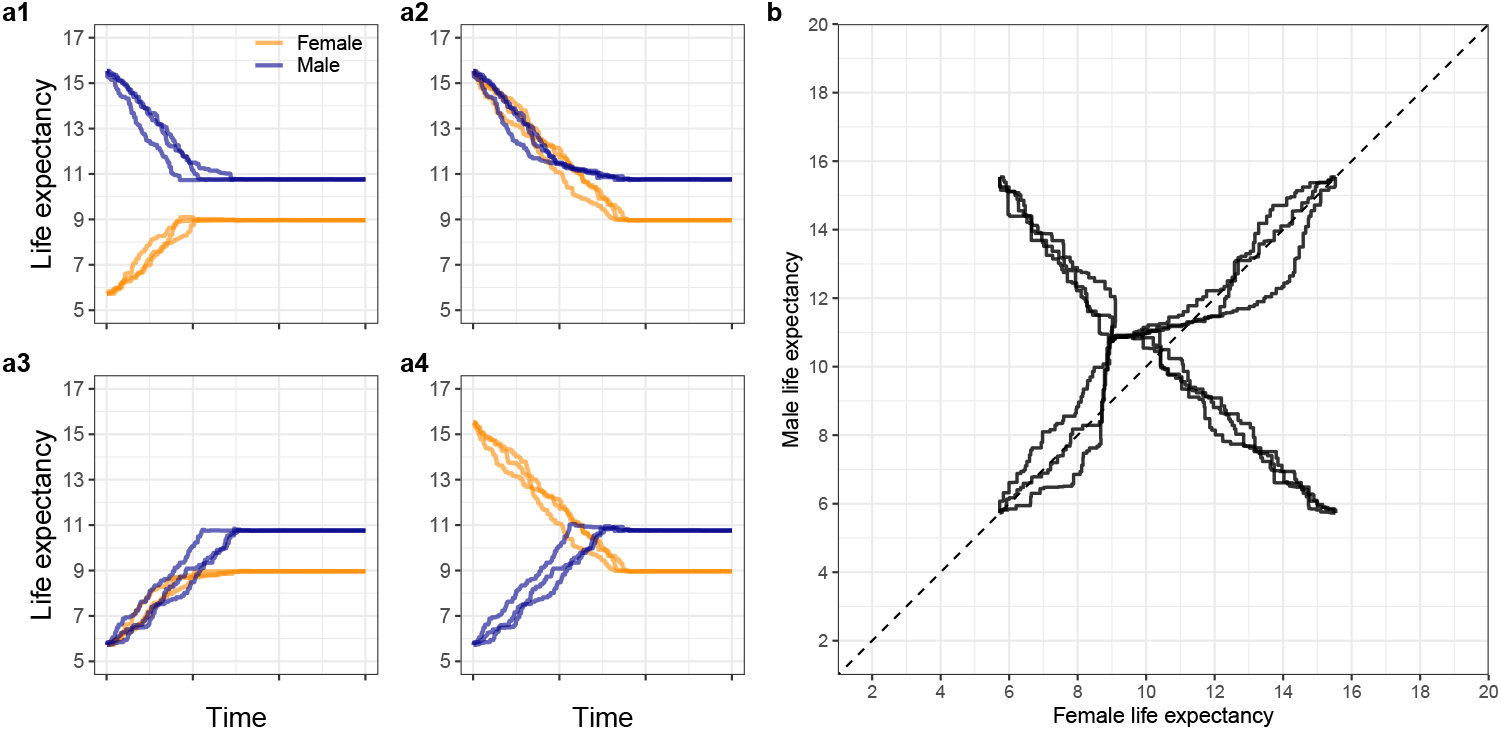
**a1-a4** Evolution of the male and female life expectancies through time towards ESS values. **B** The same data as in **a** but plotted through phenotype space instead of time.

### Effect of mating efficiency

Whilst most mating functions (e.g., a minimum function or harmonic mean) assume that all available individuals in the population will mate under balanced sex ratios, the minharmonic mating function [31] that we implement in this model allows for arguably more realistic scenarios in which not all individuals may mate despite sex ratios being balanced. This is achieved by setting the mating efficiency parameter (*η*) to values less than 1.0. In the main manuscript, we present results for *η* = 0.8 (*i*.*e*., 80% of the available single individuals can form unions under balanced sex ratios), but we also investigated the effect of varying this parameter (Fig. S2). Our main conclusions are robust to variations in mating efficiency, yet more efficient mating slightly reduces life expectancy in both sexes.

**Fig. S2:**
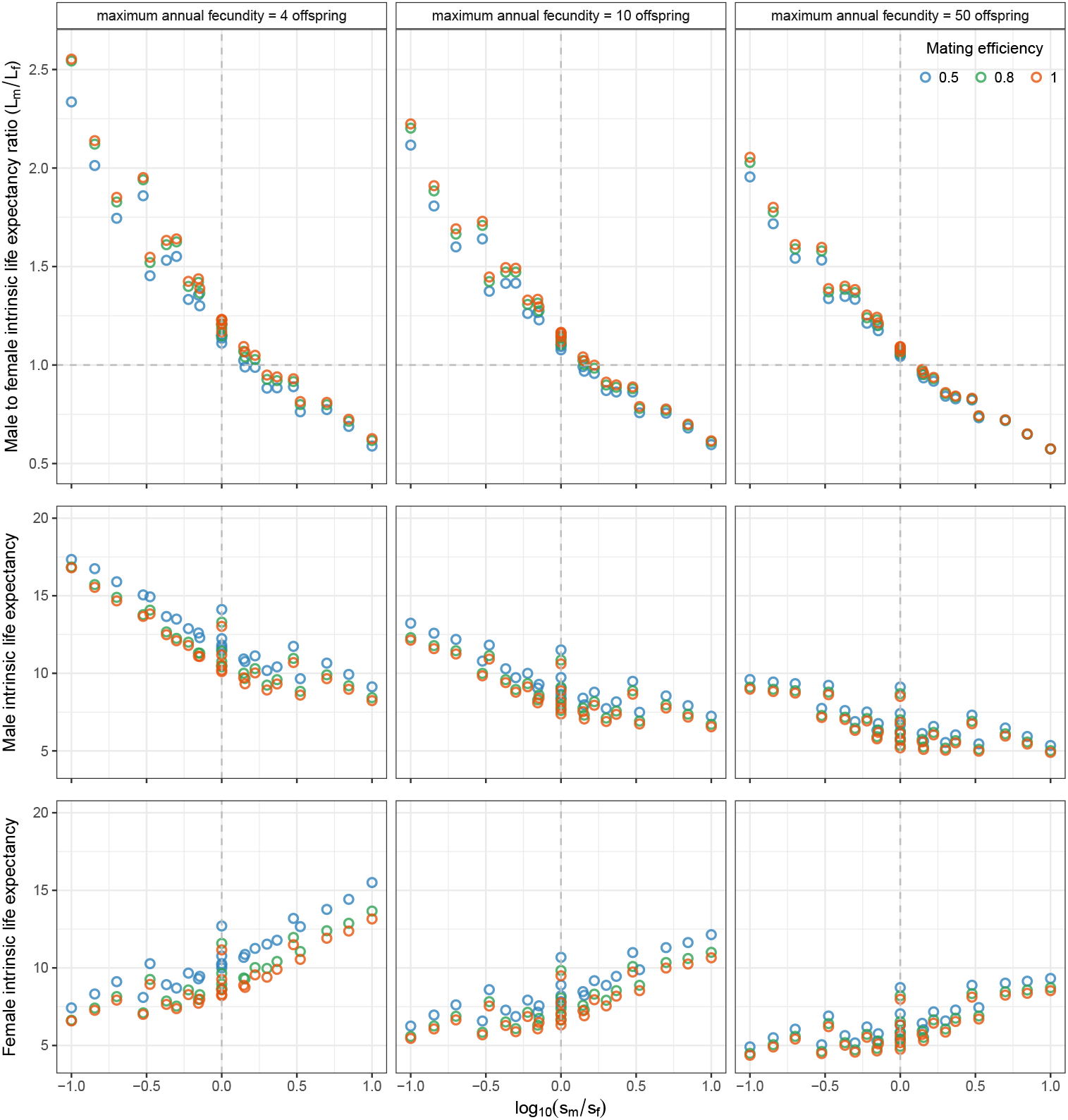
Male-to-female intrinsic life expectancy ratio (top), and the male (middle) and female (bottom) intrinsic life expectancies under various combinations of relative trade-off strengths in each sex, across varied maximum annual fecundity values (columns) and mating efficiency values (point colours). Figure conventions as in Fig. 3.

### Age-specific fertility across all trade-off strengths

**Fig. S3:**
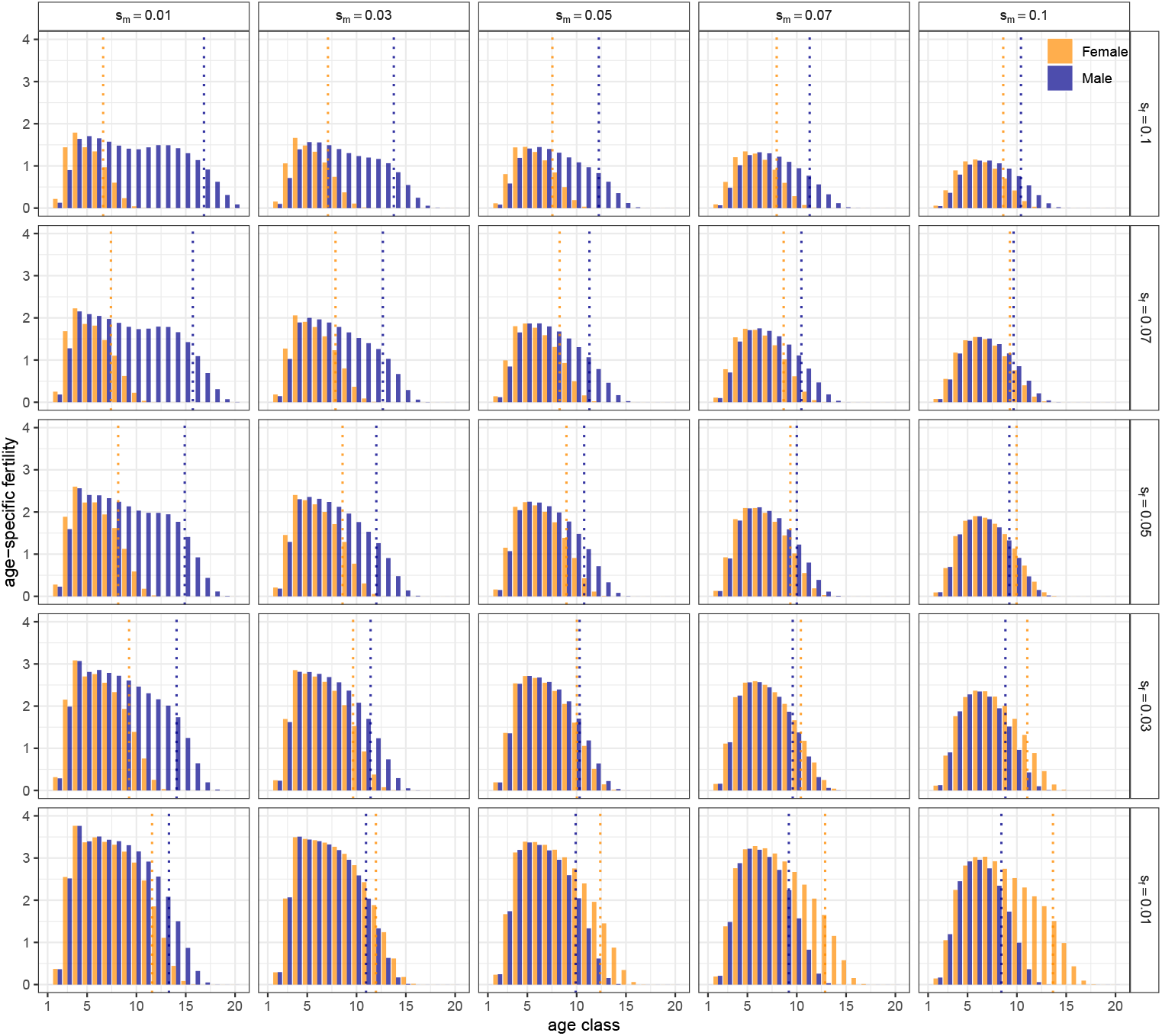
Age specific-fertility (the average number of offspring produced at a given age, conditioned on survival to that age) of males and females across the range of sex-specific trade-off combinations. Dotted vertical lines represent the life expectancy at adulthood of each sex.

### Details of matrix model construction

### Resident dynamics

#### Population vector

The model consists of a population of single males, single females, and monogamous unions (consisting of one male and one female), with each individual being of age class 1 to *ω*. The population state at time *t* can therefore be described by the following *ω* × (2 + *ω*) matrix:

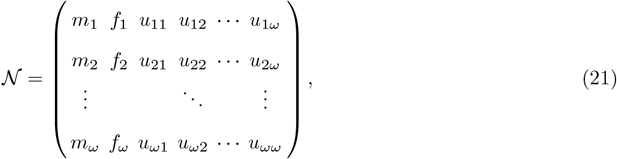

where *m*_*x*_ and *f*_*x*_ represent the number of males and females of age *x*, respectively, and *u*_*xy*_ represents the number of unions between females of age *x* and males of age *y*. This two-dimensional array is transformed into a population state vector using the vec operator, which stacks the columns on top of each other:

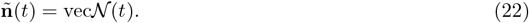

This population state vector consists of the following blocks:

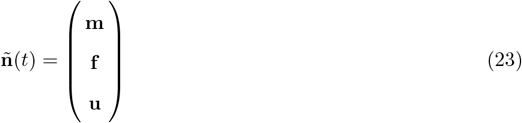

where

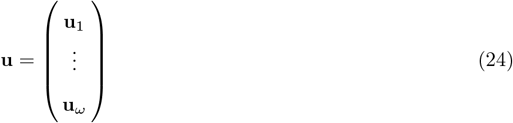

and **u**_*x*_ contains all the unions with females of age *x*. Specifically, with the age classes explicit, the sub-vectors are given by:

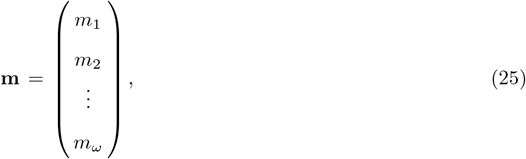

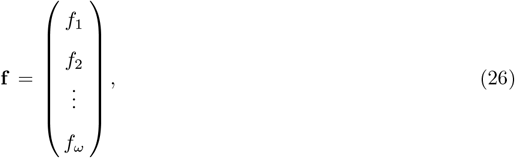

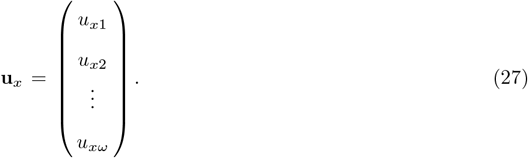

### Projection matrix

Given the three blocks of the population vector (23), the population projection matrix has following 3 × 3 structure:

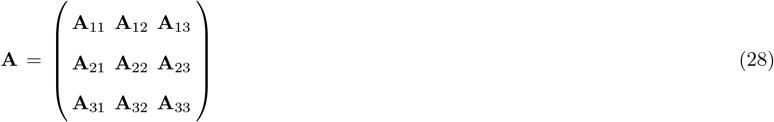

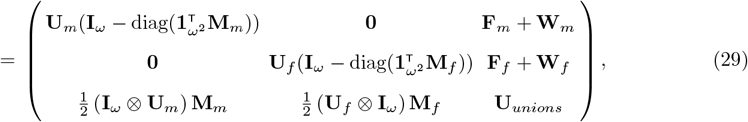

where **U**_*m*_ and **U**_*f*_ describe the age-specific survival probabilities of males and females, respectively, and **M**_*m*_ and **M**_*f*_ contain the age-specific mating probabilities (*i*.*e*., transition into unions) respectively for males and females. **U**_*unions*_ contains the age-combination-specific survival rates of unions, **F**_*m*_ and **F**_*f*_ contain the reproductive rates of unions to produce sons and daughters, respectively, and **W**_*m*_ and **W**_*f*_ describe the widowing rates of unions to produce single male and females, respectively. There are four main processes that are described in the above 3×3 projection matrix: remaining as a single (**A**_11_ for single males and **A**_22_ for single females), singles transitioning to unions (**A**_31_ for males and **A**_32_ for females), remaining as a union (**A**_33_), and single production from unions through reproduction and widowing (**A**_13_ for males and **A**_23_ for females). Each process and its relevant sub-matrices are described in more detail below.

### Remaining as a single

A single individual remains in this stage if they survive to the next age class and do not mate (transition to a union). That is, we multiply the survival probability of a given individual with one minus the probability that it mates.

Age-specific survival probabilities are described in matrix **U**_*m*_ for males and **U**_*f*_ for females. Considering an illustrative example of *ω* = 3, these matrices will have the following structures:

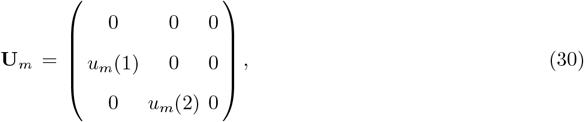

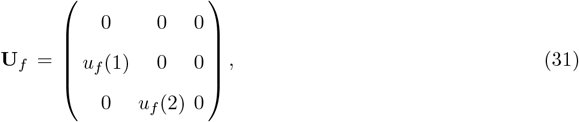

where *u*_*i*_(*x*) gives the probability that an individual of sex *i* survives from age class *x* to age class *x* + 1:

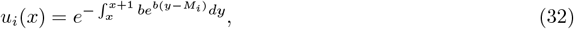

where *b* is the rate of increase of mortality with age and *M*_*i*_ is age at onset of senescence for individuals of sex *i*, and as such 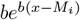 is the Gompertz hazard rate [57]. Note that individuals die when they reach the maximum age class *ω*, and thus *u*_*i*_(*ω*) = 0.

To calculate an individual’s mating probability, we use a flexible, frequency-dependent *minharmonic mating function* [31]. We first calculate the size of the mating pool. The number of single males available to mate is given by:

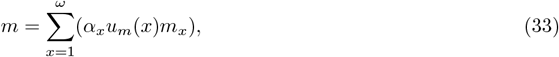

and likewise, the number of single available females is:

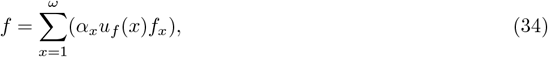

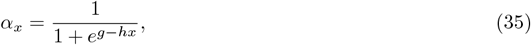

where *g* and *h* determine the shape of this logistic maturation function. As age *x* increases, *α*_*x*_ approaches 1, describing a gradual transition from reproductive immaturity to maturity. The total size of the mating pool is:

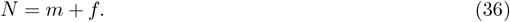

By summing over all age classes, we assume that mating is random with respect to age among individuals that are sufficiently mature to mate (as defined by equation (35)). Following [31], the expected number of unions that form each time step is given by the minharmonic mating function:

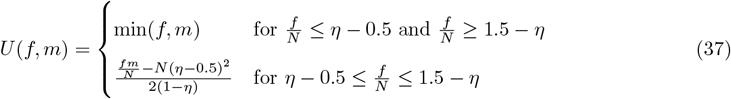

where *η* is the mating efficiency; the probability of mating at balanced operational sex ratios. For example, *η* = 1 means all available individuals form unions at balanced sex ratios, whereas *η* = 0.8 implies that only 80% of the mating pool does so, despite the balanced sex ratio. This mating function ensures that the number of unions formed is limited by the rarer sex under skewed sex ratios, and approximates the harmonic mean between sexes under balanced sex ratios. We can then use this to calculate the mating probabilities for specific union age combinations. For *ω* = 3 age classes the male mating matrix has the structure:

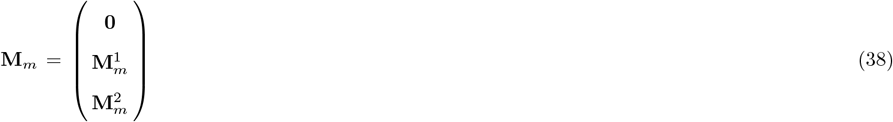

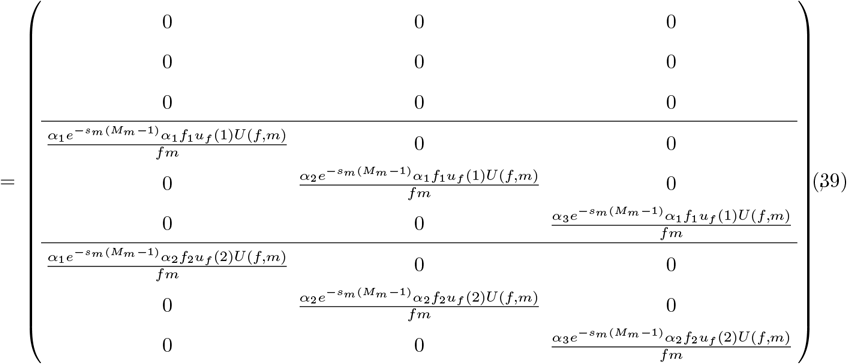

where 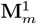 contains transitions of single males into unions with females that are currently age 1 but survive to age into age class 2. More generally for males aged *y* mating with females aged *x*:

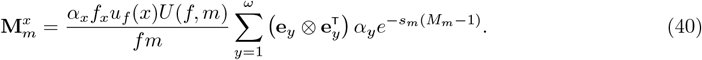

Similarly, for *ω* = 3 age classes the female mating matrix has the structure:

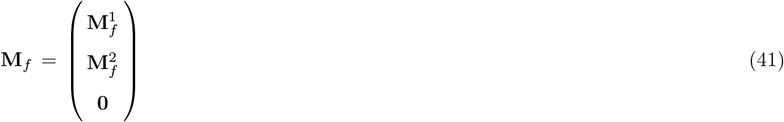

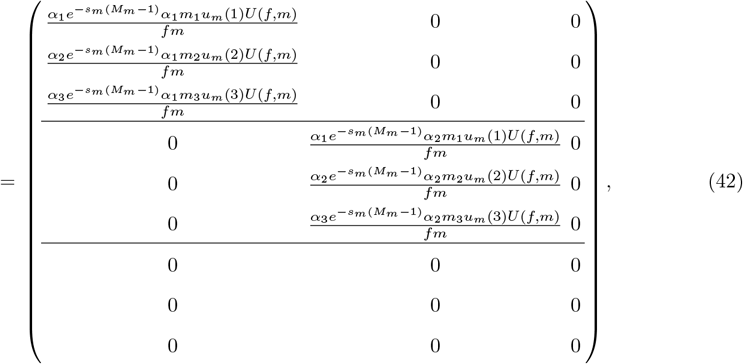

where 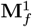 contains transitions of single females into unions who are currently aged 1 but will survive and age into age class 2. More generally for females aged *x* mating with males aged *y*:

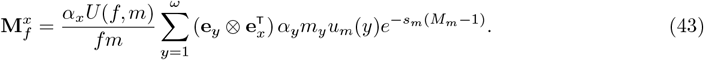

In summary, individuals stay single provided they survive and do not mate to form a union. The relevant elements of the projection matrix (equation (28)) therefore take the form 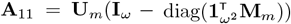 for single males and 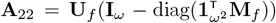 for single females, where **I**_*ω*_ is an *ω* × *ω* identity matrix containing only zeroes except for 1s across the diagonal, and 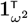 is 1 × *ω*^2^ row vector of 1s. The multiplication 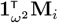 creates a vector with the column sums of the mating matrix for an individual of sex *i*, thus giving the total mating probability for each age class, and we use ‘diag()’ to produce a matrix with these probabilities across the diagonal. Since we want the probability that an individual *does not* mate, we need to calculate 1 minus the mating probability, which is achieved using 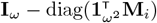. We can then multiply these “not-mating” probabilities with the age-specific survival probabilities in **U**_*m*_ or **U**_*f*_ to give the total probability that an individual stays as a single.

### Singles transitioning to unions

Singles transition to unions provided that they survive and find a mate (*i*.*e*., we multiply the probability of survival with the probability of mating). The relevant elements of the projection matrix (equation (28)) therefore take the form 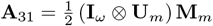 for males and 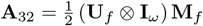 for females, where **I**_*ω*_ is an *ω* × *ω* identity matrix containing only zeroes except for 1s across the diagonal. Given the structure of the male mating probabilities within **M**_*m*_ (equation (38)) where male age varies across the diagonal within each block, we use (**I**_*ω*_ ⊗ **U**_*m*_) to repeatedly place **U**_*m*_ across the diagonal of the resulting block matrix (such that male age varies across the diagonal within each block) before it can be multiplied with **M**_*m*_. Given the structure of female mating probabilities within **M**_*f*_ (equation (41)) where female age varies between blocks, we use (**U**_*f*_ ⊗ **I**_*ω*_) to group each element of **U**_*f*_ and expand them into a block before moving to the next element (such that female age varies between blocks) before it is multiplied with **M**_*f*_. Finally, given that a single individual resident constitutes one half of a monogamous resident-resident union, the union formation rates in the projection matrix for each sex are multiplied by a factor of 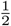.

### Remaining as a union

Unions survive to the next time step, provided both the male and female members of them survive. In this case, we also wish to move both members of the union up one age class. We achieve this using a vec-permutation framework, in which we define

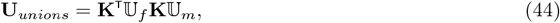

where **K** is a vec-permutation matrix (see [69] for derivation), and for *ω* = 3 age classes:

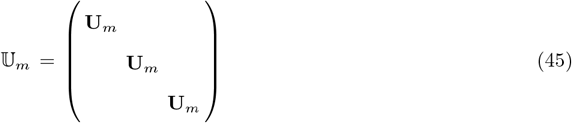

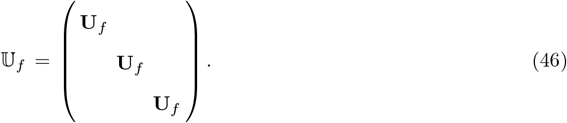

In this way, (reading equation (44) from right to left) we first consider the survival of males in each union, before re-arranging the population vector using **K** to then consider the survival of females, before using **K**^T^ to return the population vector to its original structure. Another way to visualise this process is by substituting equations (45) and (46) into equation (44) to give the matrix structure:

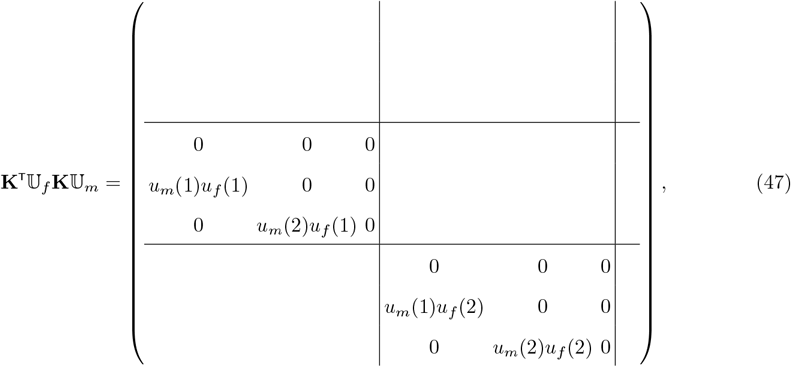

where the empty blocks are all filled with zeroes. This matrix has the block structure of the survival part of a Leslie matrix, and in addition the two non-empty blocks are themselves Leslie matrices. The blocks move vectors of unions within a specific female age to the next female age class, and then within the blocks we move unions with a specific male age class to the next age class.

### Single production from unions

Singles are produced from unions in two ways. First, unions can reproduce to create new single male and female individuals of age class 1. We describe this process in the fecundity matrices **F**_*m*_ and **F**_*f*_, respectively. Second, widowing can occur when one union member dies, and the remaining surviving partner is returned to the single population to potentially re-mate. We describe this process in the matrices **W**_*m*_ for widowed males and **W**_*f*_ for widowed females.

The fertility matrices **F**_*m*_ and **F**_*f*_ contain reproductive rates per union, which are the same for all combinations of parental ages. Offspring produced are of the first age class, and unions containing individuals of age 1 do not exist (mating to form a union is one time step before reproducing). So for *ω* = 3:

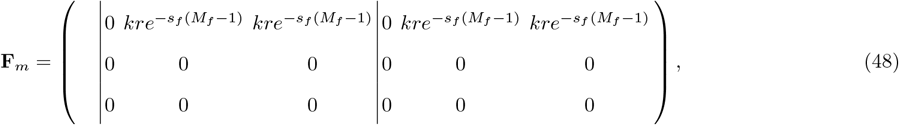

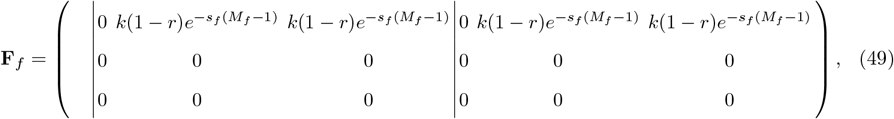

where *k* is maximum annual union fecundity and *r* is the primary sex ratio (proportion of male offspring).

Males in unions become widowed if they survive but their female partner does not. The matrix that produces single males from unions, for *ω* = 3, has the structure:

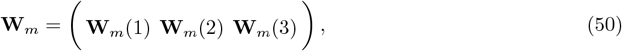

where **W**_*m*_(*x*) contains the widowing rates of males in unions with females of age *x*:

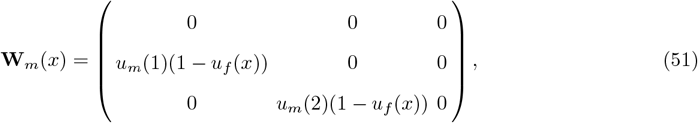

but males of age 1 cannot be mated yet, and so there are no unions with age 1 males and therefore the first non-zero entry in this matrix will never get used, or will always act on a zero entry of the population vector. We can thus start the sum in the general formula for **W**_*m*_(*x*) from *y* = 2:

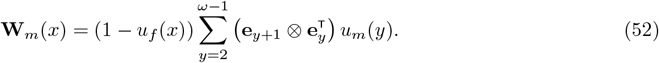

Similarly, females become widowed when they survive but their male partner does not. The matrix that produces single females from unions, for *ω* = 3, has the structure:

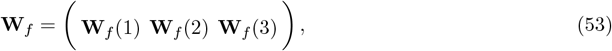

where **W**_*f*_ (*x*) contains the widowing rates of females of age *x* such that

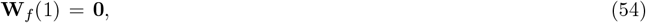

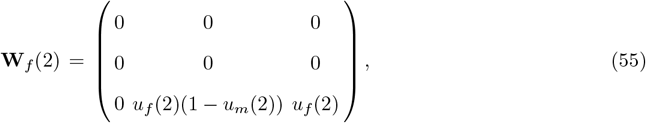

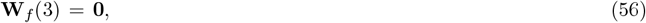

given that females aged 1 cannot be widowed as they cannot yet be mated, and females aged *ω* (3 in this example) are not widowed since they die themselves. More generally,

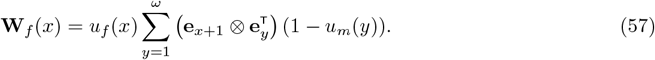

In summary, single males and females are produced from unions either as offspring or as widowed adults, such that **A**_13_ = **F**_*m*_ + **W**_*m*_ and **A**_23_ = **F**_*f*_ + **W**_*f*_ in the projection matrix (equation (28)).

### Mutant dynamics

#### Population vector

Unlike under resident dynamics, for the mutant population there are two types of union possible: resident females with mutant males, and mutant females with resident males [60]. Under an adaptive dynamics framework, we assume that the mutant is rare and thus that mutant-mutant unions have negligible effects on population dynamics, so mutant-mutant unions are not modelled here. Note that there is no projection of the mutant population vector in the model: mutant growth rate is calculated by evaluating the mutant projection matrix at the resident equilibrium population. We nevertheless show the explicit structure of the mutant population to aid understanding of the construction of the mutant projection matrix. The population vector has the structure:

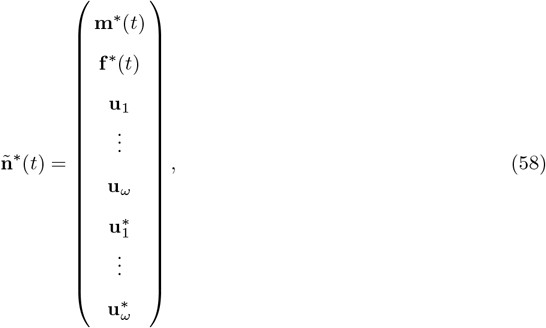

where **u**_*x*_ contains all the unions between resident females of age *x* with mutant males, and 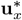 contains all the unions between mutant females of age *x* with resident males. Specifically, with the age classes explicit, the sub-vectors are given by

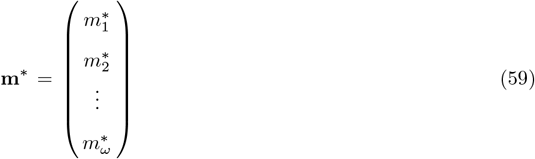

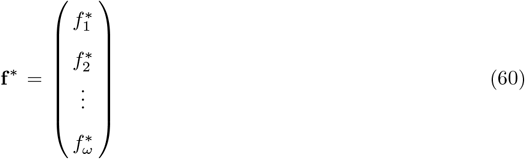

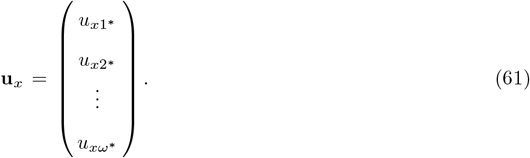

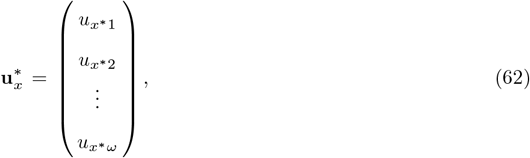

where the ∗ indicates a mutant.

#### Projection matrix

Given the structure of the mutant population vector (equation (58)), the mutant population projection matrix will have the structure:

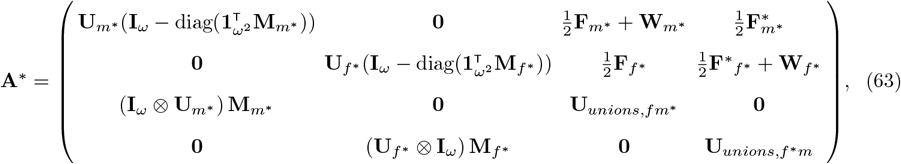

Where 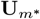 and 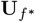 describe the age-specific survival probabilities of mutant males and females, respectively, and 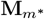 and 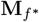 contain the age-specific mating probabilities (*i*.*e*., transition into unions) respectively for mutant males and females with a resident of the opposite sex. 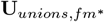 and 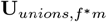 contain the age-combination-specific survival rates of the two union types. 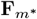 gives the reproductive rates of unions with resident females to produce mutant sons, and likewise 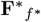 for unions with mutant females to produce mutant daughters. Note that 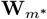 and 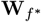 describe the widowing rates of unions with a resident of the opposite sex to produce single mutant male and females, respectively. Since we model sequential alternating mutations of *M*_*m*_ and *M*_*f*_, a mutation in the trait of one sex means mutants of the opposite sex are phenotypically residents (but still carry the mutation that can be passed on to offspring). More specifically, when *M*_*m*_ mutates to 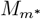, mutant females still express the resident female *M*_*f*_ value, which is thus is used in the calculations in the sub-matrices 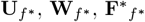 and 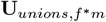 Similarly, when *M*_*f*_ mutates to 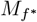, mutant males still express the resident male *M*_*m*_ value, which is thus used in the calculations in the sub-matrices 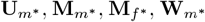 and 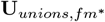.

As with resident dynamics, we have four main processes described in this mutant 4 × 4 population projection matrix: remaining as a single (A^*^ _11_ for single mutant males and A^*^_22_ for single mutant females), singles transitioning to unions (A^*^_31_ for mutant males with a resident female, and A^*^_42_ for 38 mutant females with a resident male), remaining as a union (A^*^_33_ for mutant males with a resident female, and A^*^_44_ for mutant females with a resident male), and single production from unions (A^*^_13_ for producing single mutant males and A^*^_24_ for single mutant females). Each process and its relevant sub-matrices are described in more detail below.

### Remaining as a single

As with residents, mutants remain single mutants provided they survive and do not mate. Therefore, this section of the mutant projection matrix has the same general structure of that for the residents. We define the survival matrices for mutants in the same structure:

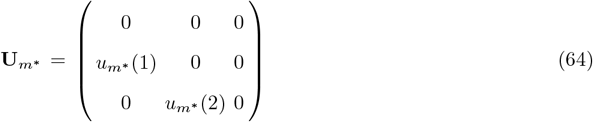

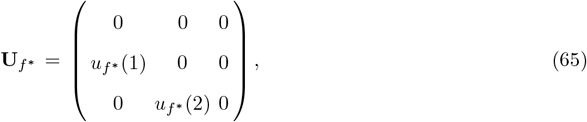

Where 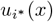 is the probability that a mutant of sex *i* survives from age *x* to age *x* + 1. The survival probability 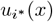 is calculated in the same way as for residents (equation (32)), but using the mutant ages of onset of senescence,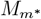 and 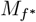, instead of those of the residents.

The marriage matrices also take a similar structure as those for the resident population. Since we assume the mutants have a negligible effect on the resident population, we still define the mating pool as consisting of residents only (*N* in equation (36); *m* in equation (33); *f* in equation (34)). We use these aspects of the resident mating pool to calculate the conditional value of *U* (*f, m*) (equation (37)). Similarly as for the residents, for *ω* = 3 age classes the mutant male mating matrix has the structure:

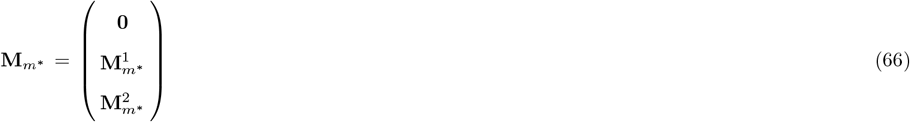

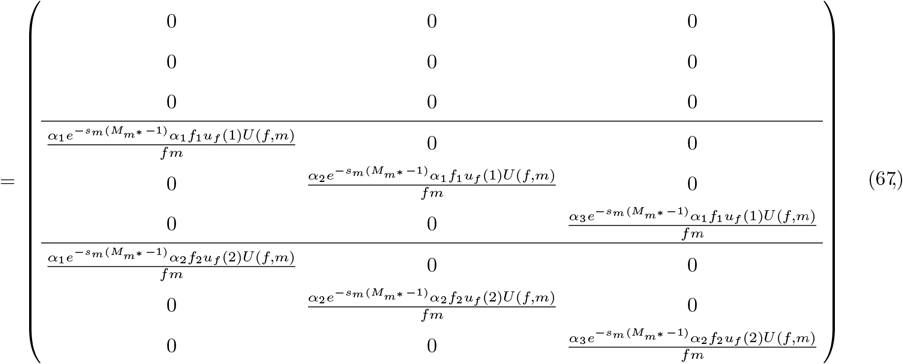

where 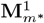 ontains transitions of single males into unions with females that are currently age 1 but survive to age into age class 2. This is identical to the male mating matrix of the residents, only here we use 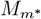 in the trade-off term instead of *M*_*m*_. More generally for mutant males aged *y* mating with resident females aged *x*:

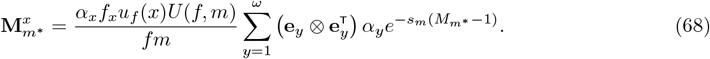

Since mutant females form unions with resident males, we use *M*_*m*_ in the trade-off term, and as such the mutant female mating matrix is identical to that of resident females (equation (43); 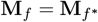).

Similarly to the residents, the relevant elements of the mutant projection matrix (equation (63)) take the form 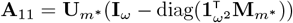 for single mutant males and 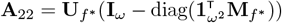 for single mutant females, since an individual stays single provided they survive and do not mate into a union (*i*.*e*., multiply the survival probability with one minus the mating probability).

#### Singles transitioning to unions

As with the residents, mutant singles also transition to unions provided that they survive and find a mate (*i*.*e*., we multiply the probability of survival with the probability of mating). The relevant elements of the projection matrix (equation (63)) therefore take the form 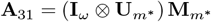 for mutant males and 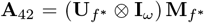 for mutant females. These terms are identical in structure as those for the residents, except we do not multiply by a factor of 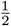 here since we are explicitly moving a single mutant individual into the union, not the resident partner.

#### Remaining as a union

Unions survive to the next time step, provided both partners within them survive. In this case, we also wish to move both partners up one age class. We achieve this in the same way as for residents (equation (44)), but use 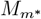 or 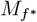 for the calculation of relevant survival probabilities. That is, we define

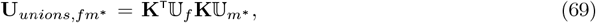

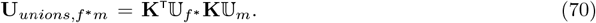

**Single production from unions**

As for with the resident population, mutant singles can also be produced from unions either as offspring through union reproduction, or as a widowed adult if the union partner does not survive. Since in mutant dynamics we only explicitly model the production of mutant offspring, 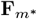 describes the production of mutant males from a union of a resident female and mutant male, and 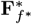 describes the production of mutant females from a union of a mutant female and resident male. For *ω* = 3:

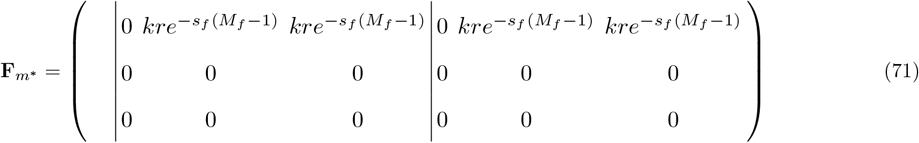

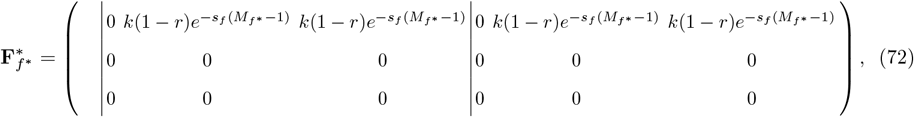

and note that since resident females are reproducing in unions of resident females and mutant males, 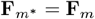 (equation (48)).

Widowing of mutants also works in the same way as for residents, only using 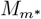 or 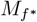 for the calculation of relevant survival probabilities. As such, the matrix that produces single mutant males from unions of resident females and mutant males, for *ω* = 3, has the structure:

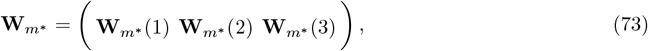

where 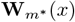 contains the widowing rates of mutant males in unions with resident females of age *x*:

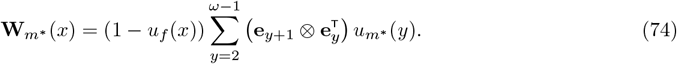

Similarly, the matrix that produces single mutant females from unions of mutant females and resident males, for *ω* = 3, has the structure:

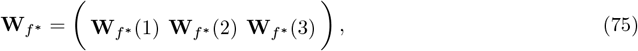

where 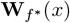 contains the widowing rates of mutant females of age *x* such that

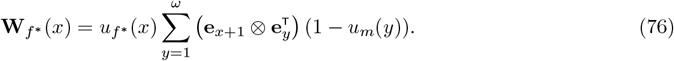

In summary, as with the residents, single mutant males and mutant females are produced from unions either as offspring or as widowed adults, such that 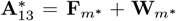 and 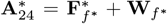 in the projection matrix (equation (63)).

